# Network activity influences the subthreshold and spiking visual responses of pyramidal neurons in a three-layer cortex

**DOI:** 10.1101/087288

**Authors:** Nathaniel C. Wright, Ralf Wessel

## Abstract

A primary goal of systems neuroscience is to understand cortical function, which typically involves studying spontaneous and sensory-evoked cortical activity. Mounting evidence suggests a strong and complex relationship between the ongoing and evoked state. To date, most work in this area has been based on spiking in populations of neurons. While advantageous in many respects, this approach is limited in scope; it records the activities of a minority of neurons, and gives no direct indication of the underlying subthreshold dynamics. Membrane potential recordings can fill these gaps in our understanding, but are difficult to obtain *in vivo*. Here, we record subthreshold cortical visual responses in the *ex vivo* turtle eye-attached whole-brain preparation, which is ideally-suited to such a study. In the absence of visual stimulation, the network is “synchronous”; neurons display network-mediated transitions between low- and high-conductance membrane potential states. The prevalence of these slow-wave transitions varies across turtles and recording sessions. Visual stimulation evokes similar high-conductance states, which are on average larger and less reliable when the ongoing state is more synchronous. Responses are muted when immediately preceded by large, spontaneous high-conductance events. Evoked spiking is sparse, highly variable across trials, and mediated by concerted synaptic inputs that are in general only very weakly correlated with inputs to nearby neurons. Together, these results highlight the multiplexed influence of the cortical network on the spontaneous and sensory-evoked activity of individual cortical neurons.

---

Spikes are fundamental to cortical function; they are the means by which individual neurons receive and transmit information, and are the unit of language for cortical ensembles that encode sensory information. Understandably, then, most studies of cortical sensory responses have focused on the spiking activities of (increasingly large) populations of neurons. One recurring theme in this vast body of literature is the rich relationship between ongoing and evoked activity. Yet the spike-based approach yields an incomplete picture of this relationship (and sensory cortex generally), for three reasons. First, it reveals the activity of a minority of neurons; most cells spike very rarely, if at all^1^, and of those that do, few have spike rates sufficient for certain analyses^2^ (Figure 1a). Second, neuronal populations defined by the recording device’s field of view are unlikely to represent complete cortical microcircuits (Figure 1b). (While the local field potential (LFP) is both easily obtained and less susceptible to the first issue, this signal too is ultimately defined by the device (Figure 1b).) Third, the purely suprathreshold view of cortex leaves certain important questions unanswered. For example, some competing hypotheses of cortical function are not easily distinguishable by the spiking statistics of small populations, but predict very different subthreshold dynamics for individual neurons^3–6^. This third point motivates recording the membrane potential, which in fact neatly addresses the first two issues as well. First, each neuron samples an enormous and biologically-relevant presynaptic pool (Figure 1b), and thus the subthreshold membrane potential communicates information about spiking in that pool (Figure 1c). Second, the membrane potential is a spike-rate independent measure of activity in the recorded neuron, and therefore gives voice to sparse-spiking neurons (Figure 1c). For these reasons, it is vital to supplement the literature on cortical spiking with studies of subthreshold sensory responses.

**Figure 1.**
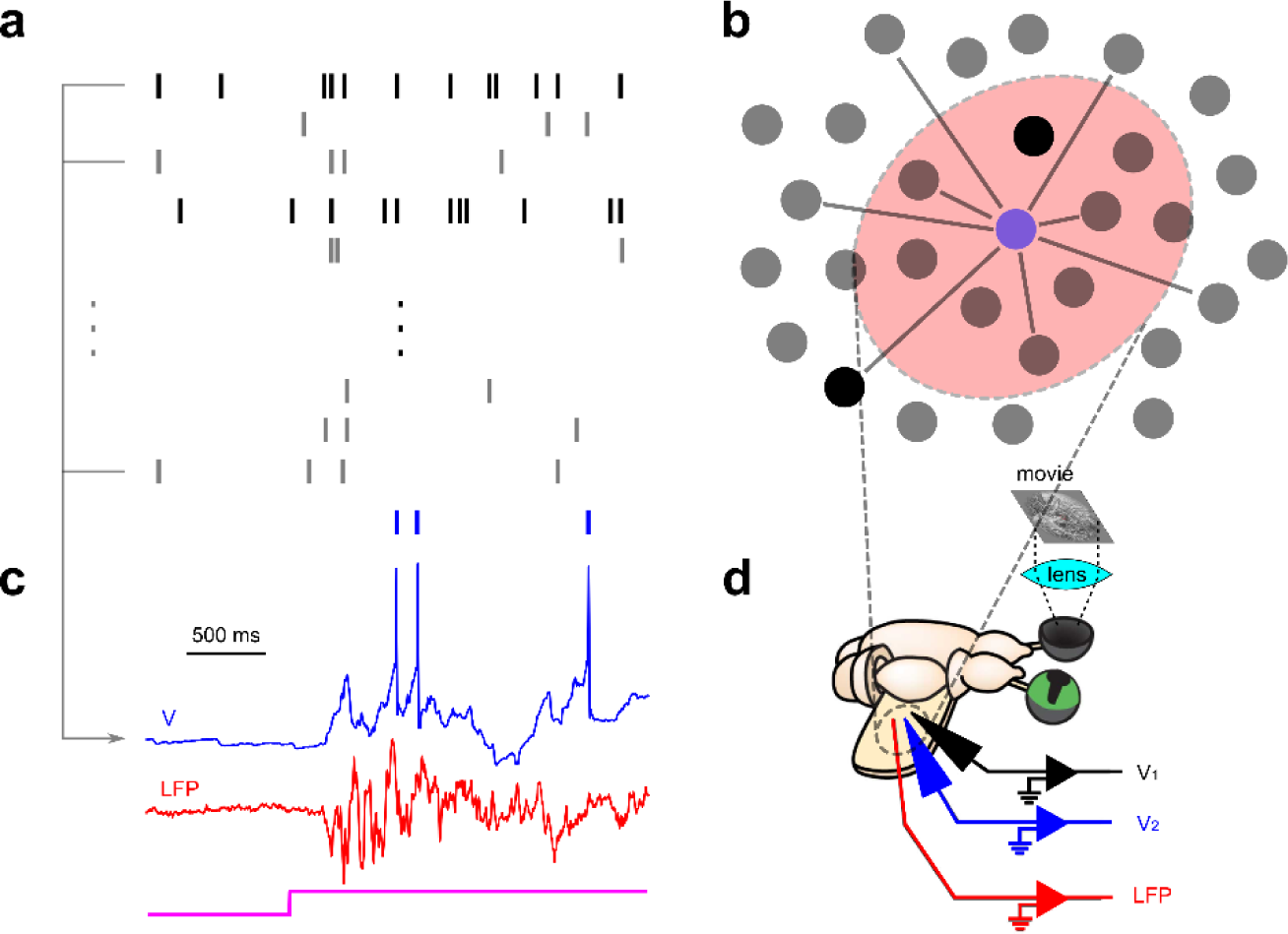
Whole-cell recordings supplement our understanding of cortical activity. (a) Ongoing and sensory evoked cortical spiking is sparse; a minority of neurons (high opacity rasters) spike often enough to give sufficient spike counts for some analyses. (b) Devices used to record population spiking activity define sets of neurons which are primarily sparse-spiking (low-opacity dots). These populations do not in general represent cortical microcircuits. The local field potential (LFP, red region) is useful for estimating synaptic activity, but only in a small electrode-defined region. A given neuron (e.g., blue dot) is a useful network sub-sampler; it receives inputs from a biologically relevant presynaptic pool spanning large regions of cortex. (c) The membrane potential of a cortical neuron (blue trace) provides (i) a spike-rate independent measure of neuronal activity, and (ii) estimates the spiking activity of the presynaptic pool. The simultaneously-recorded nearby LFP (red trace) is useful for interpreting this subthreshold activity in the context of local synaptic activity. (d) We simultaneously record the membrane potentials from groups of cortical neurons (as well as the nearby LFP) in the turtle eye-attached whole-brain *ex vivo* preparation, during ongoing and visually-evoked activity.

This is easier said than done; while stable patch clamp recordings are readily achievable in slice and cell culture, they are extremely difficult to obtain *in vivo*. Consequently, studies of subthreshold visual responses are relatively rare. Accordingly, we have recorded ongoing and visually-evoked subthreshold and spiking activity from cortical pyramidal neurons in the *ex vivo* turtle eye-attached wholebrain preparation (Figure 1d), which is ideally-suited to such an investigation^7,8^. Specifically, it allows for stable patch clamp recordings (lasting up to two hours) from neurons in a three-layer visual cortex (analogous to mammalian piriform cortex and hippocampus^9–11^) subject to inputs from an intact visual pathway. In some cases, we simultaneously record the nearby LFP to interpret the subthreshold events in the context of local synaptic activity (Figure 1d).

Here, we present four key observations. First, we find that the ongoing network state is “synchronous”; subthreshold activity reveals a relatively quiescent “low-conductance” state that is frequently interrupted by “high-conductance” events. The increase in synaptic conductance results in slow-wave activity (or broad membrane potential depolarizations) with fast nested subthreshold fluctuations. High-conductance event onsets are correlated across pairs of nearby neurons, and coincide with oscillations in the nearby local field potential (suggesting they are network-generated). Second, brief and extended visual stimulation evoke persistent high-conductance states that are qualitatively similar to larger spontaneous synaptic events, and less synchronous than pre-stimulus activity. Spiking in this state is sparse and highly variable across trials (in terms of both precise timing and spike counts). Third, visual stimulation that interrupts or follows soon after large, spontaneous events evokes responses that are muted relative to the average. Finally, while the evoked state is asynchronous at long time scales, spikes are preceded by concerted excitatory synaptic inputs that are not in general coordinated across neighboring neurons. These brief, pre-spike depolarizations are less pronounced for neurons exhibiting stronger slow-wave fluctuations during ongoing activity.

Taken together, these results provide a rare view into the subthreshold dynamics of cortical visual responses. They highlight the effects of network-mediated synaptic activity on the spiking of individual neurons, and demonstrate the utility of the membrane potential as a tool for sampling the state of a presynaptic population. Ultimately, this diagnostic tool suggests strong network influences on single-neuron activity across multiple spatiotemporal scales.

## Results

In order to investigate the nature of spontaneous and evoked subthreshold cortical activity, we obtained whole-cell recordings from neurons in the *ex vivo* turtle eye-attached whole-brain preparation (Figure 1), both in the absence of visual stimulation (Figure 2a), and in response to brief and extended visual stimulation (Figure 3a, see Methods).

**Figure 2.**
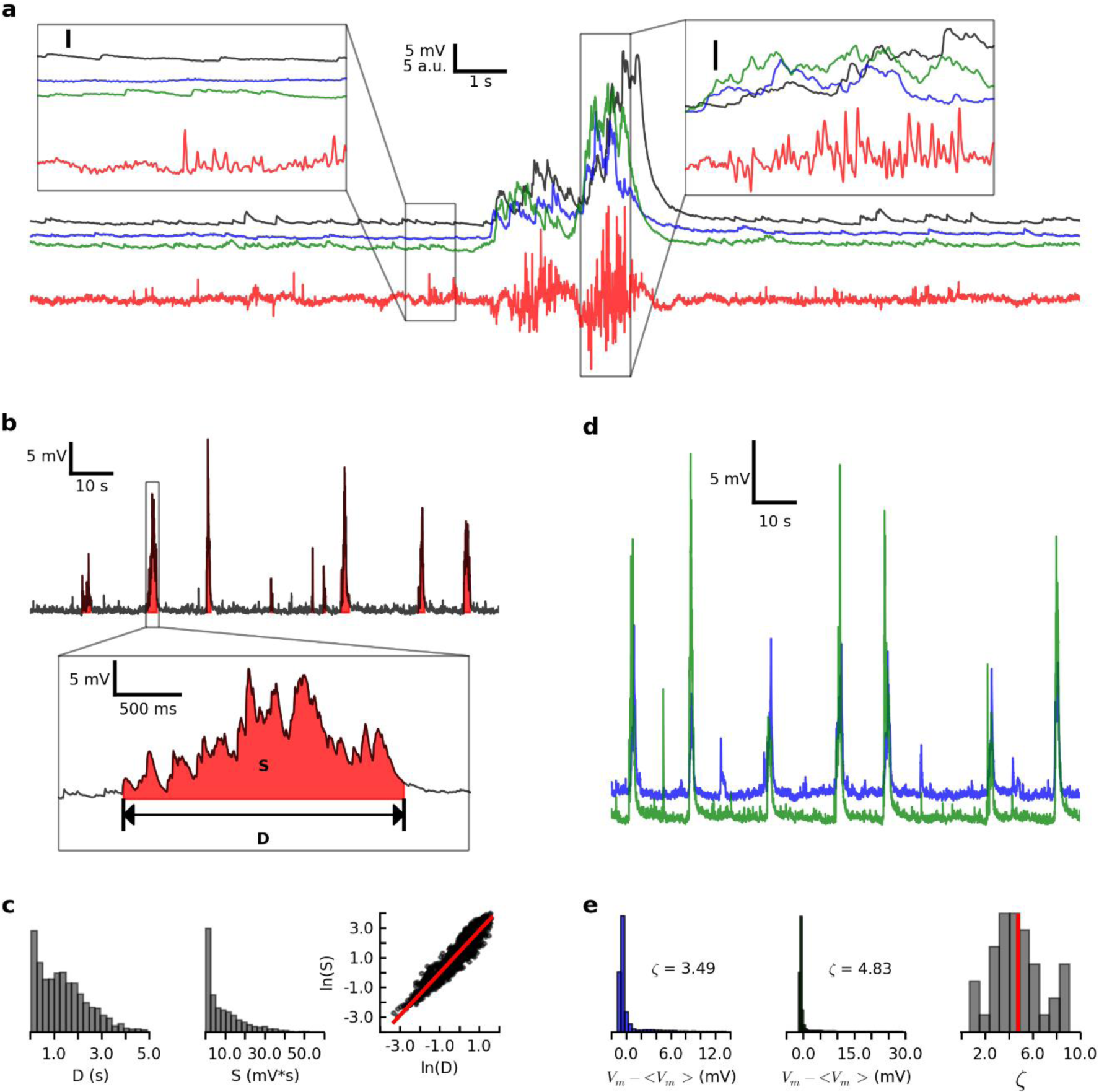
During spontaneous activity, the low-conductance membrane potential state is interrupted by broadly-correlated high-conductance events. (a) Subthreshold membrane potentials from three simultaneously-recorded neurons (black, blue, and green traces) and the nearby LFP (red trace) in the absence of visual stimulation. Left (right) inset: 1 s of “low-conductance” (“high-conductance”) activity. (b) Top: spontaneous membrane potential recording, with high conductance events filled in red. Bottom: enlarged view of an individual spontaneous high-conductance event. D is event duration, and S (area under curve) is event size. (c) Distributions of high-conductance event durations (left) and sizes (middle), for 1389 events from 40 cells in 16 turtles. Right: natural log of size vs. natural log of duration, for all events. Red line indicates significant linear regression fit (r = 0.95, slope = 1.43, P < 1 × 10^−300^). (d) Spontaneous subthreshold membrane potentials from two simultaneously-recorded neurons. (e) Left and center: distributions of mean-subtracted membrane potentials for the two neurons in (e), with distribution skew (ζ). Right: distribution of skews for spontaneous activity for 40 neurons from 16 turtles. Red line indicates across-cell mean skew.

**Figure 3.**
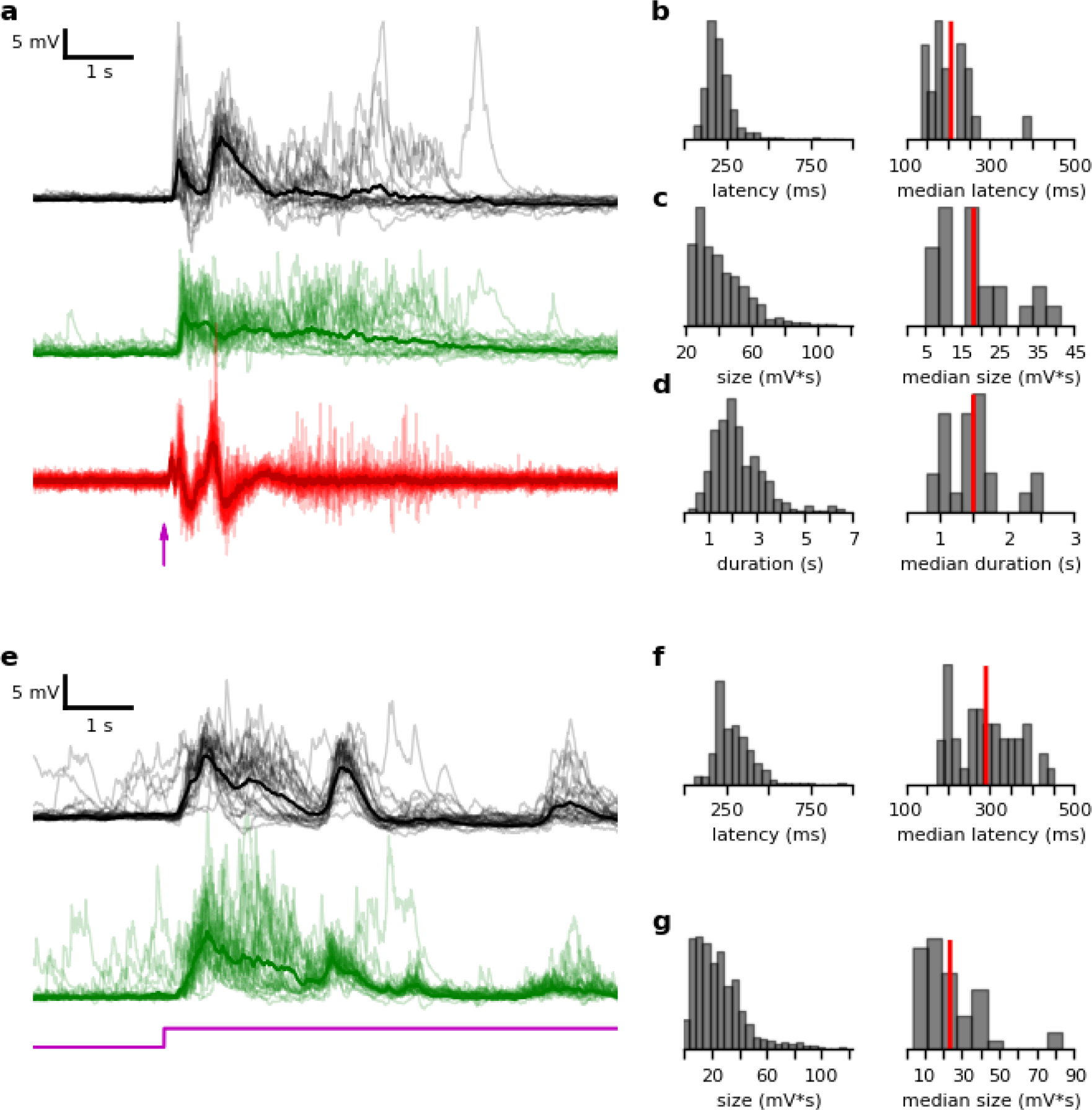
Subthreshold visually-evoked activity is highly variable across cells and trials. (a) Membrane potentials from two simultaneously-recorded neurons (black and green traces) and the nearby LFP (red traces), with individual trials in low opacity, and across-trial averages in high opacity. Stimulus is whole-field red flash (flash onset at arrow, see Methods). (b) Left: distribution of subthreshold response latencies (see Methods) for 559 trials from 23 cells in 9 turtles. Stimulus is either red whole-field or sub-field flash (see Methods) Right: distribution of across-trial median response latencies for these cells. Red line indicates across-cell average. (c) Same as in (a), but for response size (see Methods). (d) Same as in (a), but for response duration (see Methods). (e) Same as in (a), but stimulus is naturalistic movie (see Methods), for a different pair of simultaneously-recorded neurons. (f, g) Same as in (b, c), but for extended visual stimuli (1087 trials from 48 cells in 15 turtles, see Methods).

### Spontaneous activity is characterized by transitions between low- and high-conductance states

To help identify the effects of visual stimulation on subthreshold activity, we first sought to characterize spontaneous activity. To this end, we recorded from 40 neurons while maintaining the preparation in complete darkness. For 25 of these cells, we simultaneously recorded the nearby LFP. In the absence of visual stimulation, membrane potentials were typically far from action potential threshold, displaying small, but frequent postsynaptic potentials (PSPs, Figure 2a, **left inset**). Occasionally, coordinated barrages of PSPs interrupted these periods of relative quiescence (Figure 2a). For most cells, the longer-duration barrages resulted in broad membrane potential depolarizations, with nested higher-frequency fluctuations (Figure 2a, **right inset**). The onset of large “high-conductance events” was correlated across pairs of nearby neurons, and coincided with the onset of oscillations in the nearby LFP (Figure 2a).

We used an algorithm to detect these spontaneous events (see Methods) and quantified the duration (D) and size (S) of each (considering only those between 100 ms and 5 s in duration, Figure 2b). Both S and D varied considerably across events (Figure 2c), and had a strong log-log relationship (r = 0.94, slope = 1.43, P < 1 × 10^-300^, linear regression fit, for 1362 high-conductance events from 40 cells in 16 turtles, Figure 2c, **right**).

We next sought to understand these subthreshold dynamics in the context of network activity. To do this, we made use of the fact that membrane potential distributions carry statistical signatures of the presynaptic network state. In the so-called “asynchronous” state, for example, random synaptic inputs result in subthreshold membrane potentials that evolve according to a “random walk”, and membrane potential distributions are thus approximately Gaussian (with small or negative skew^3,4^). In contrast to this scenario, we found that the largest of the spontaneous high-conductance events observed here had greater amplitudes and durations than those predicted for an asynchronous network with the same mean activity level, as captured by the long, depolarized tails of membrane potential distributions (Figure 2e, **left and center**). These tails yielded positive distribution skews (ζ) for all cells (population-average skew <ζ> = 4.74 +/− 2.08, mean +/− s.e.m., Figure 2e, **right**). As such, ongoing activity in this preparation was consistent with the well-characterized “synchronous network state” observed across a variety of mammalian cortical areas^3,4,12–16^. In this state, the subthreshold activities of individual neurons, which provide a measure of presynaptic network activity, indicate brief periods of elevated activity that are broadly coordinated across time and cortical space.

### Visual stimulation evokes high-conductance states with large across-trial variability and sparse spiking

Next, we characterized cortical responses to visual stimulation. For 23 cells from nine turtles, we recorded ongoing activity and responses to either whole-field or sub-field flashes (and obtained at least 12 valid trials, see Methods). In response to flashes, neurons received persistent barrages of synaptic inputs that were coincident with nearby LFP oscillations (Figure 3a). This activity was qualitatively similar to longer-duration spontaneous high-conductance events (compare to Figure 2a). For individual neurons, the response time course was highly variable across trials (Figure 3a). Across all neurons and trials, this was also true for response latency (Figure 3b), size (Figure 3c), and duration (Figure 3d). For this type of stimulus, evoked activity in retinal ganglion cells was unlikely to continue beyond a few hundred milliseconds after stimulus onset^17^. Thus, the late response phase was likely due to persistent intracortical and/or thalamocortical activity.

We next investigated the effects of persistent sensory input by recording from 48 cells from 15 turtles while presenting extended visual stimulation (with at least 12 valid trials, see Methods). These stimuli evoked subthreshold responses that were qualitatively similar to flash responses in the early response phase, but displayed clear temporal structure hundreds of milliseconds later (Figure 3e). Thus, although intracortical inputs were likely extremely strong in the late response phase (as suggested by the persistent responses to brief flashes), the modulatory effects of sensory input were also evident.

Finally, we characterized evoked spiking activity. Spikes were sparse in general, and highly variable across trials in terms of precise spike timing and total spike counts (Figure 4a). The population-average rate was 0.37 +/− 0.74 Hz (mean +/− s.e.m.) in the two seconds after stimulus onset (for all stimuli, N = 79 cells, Figure 4a, **inset**). Of the 79 recorded cells (with at least 12 valid trials, see Methods), 24 (or 30%) did not spike at all in this window. Of those that did, the average rate was 0.53 +/− 0.83 Hz.

**Figure 4.**
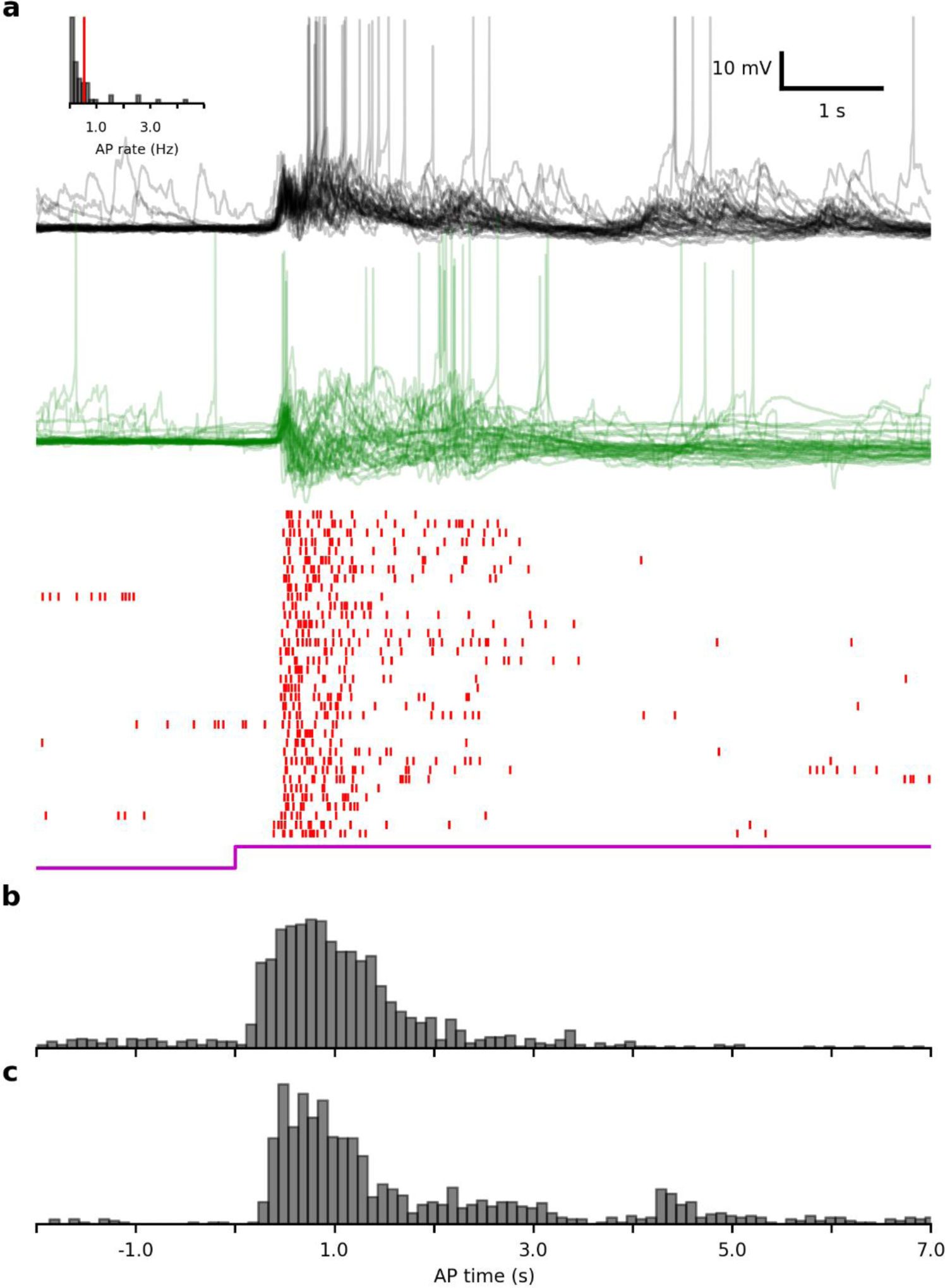
Visually-evoked spiking is sparse, and highly variable across cells and trials. (a) Membrane potentials of two simultaneously-recorded neurons (black and green traces) and nearby multi-unit activity (red rasters) during ongoing and visually-evoked activity, across multiple stimulus presentations. Rasters are stacked in ascending trial order. Inset: distribution of across-trial average evoked spike rates for 27 cells in 8 turtles, for all stimuli (see methods). (b) Peristimulus time histogram of intracellularly-recorded spikes for responses to flashes (1196 spikes from 27 cells in 8 turtles, see Methods). (c) Same as in (b), but for responses to extended stimuli (1185 spikes from 33 cells in 14 turtles, see Methods).

We inspected the time course of evoked spiking by constructing peri-stimulus time histograms (PSTHs) for all cells with at least one evoked spike in the 7 s after stimulus onset, across all trials. In response to brief flashes, most spikes occurred in the 2 s after stimulus onset (Figure 4b). Between 4 s and 7 s after stimulus onset, spike rates fell below pre-stimulus levels. That is, the strong bouts of evoked activity in the 4 s after the onset of brief flashes appeared to suppress spontaneous spiking. The early (0 to 2 s) spiking responses to movies were similar to those for flashes, but rates were elevated above those for brief flashes in the later response (Figure 4c).

### Evoked high-conductance states are broadly asynchronous

We next sought to more carefully characterize the effects of visual stimulation on the network state, as communicated by single-neuron subthreshold membrane potentials. As shown above, at broad time scales (on the order of seconds), the network was in a synchronous state during spontaneous activity; rather than hovering near action potential threshold, neurons were driven near threshold by barrages of synaptic inputs (Figure 2a). On shorter times scales (hundreds of milliseconds), evoked activity appeared closer to the description of asynchrony (Figure 3a, e). Was this, in fact, the case? That is, did visual stimulation cause a shift from synchrony to asynchrony in the network?

We addressed this question by calculating skews for residual membrane potentials (membrane potential time series with across-trial average time series subtracted) during ongoing and visually-evoked activity, for responses to all stimuli (see Methods). For each cell, we considered an “ongoing” window of pre-stimulus activity (2 s to 0 s before stimulus onset), and an “evoked” window of activity (starting at response onset, and lasting 2 s, Figure 5a, see Methods). Consistent with our observations of long spontaneous recordings (Figure 2e), ongoing skew values (Figure 5b, **top**) were typically large and positive (population-average ongoing skew <ζ> = 1.82 +/− 1.43, Figure 5c). For most cells, skew decreased from the ongoing to evoked window (Figure 5b, **bottom**), an effect that was significant for the population as a whole (evoked <ζ> = 1.14 +/− 0.67, P = 3.2 × 10^−4^ for ongoing-evoked comparison, Wilcoxon signed-rank test, Figure 5c). Thus, evoked activity was less synchronous than ongoing, on time scales of hundreds of milliseconds to seconds.

**Figure 5.**
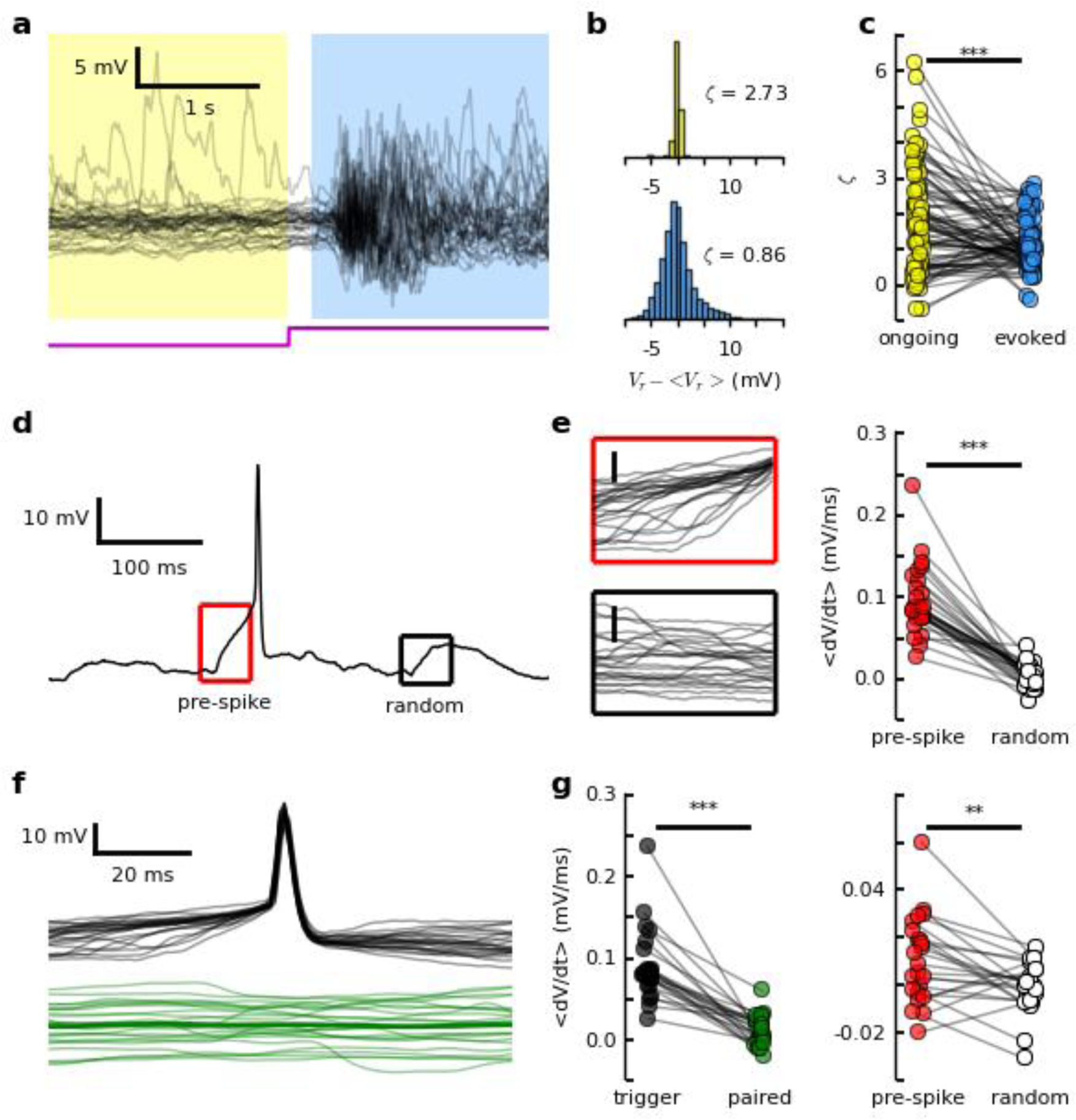
Visual stimulation affects coordination at multiple spatiotemporal scales. (a) Residual membrane potentials (see Methods) for ongoing (yellow) and evoked (blue) activity, for repeated presentations of motion-enhanced movie (see Methods). (b) Distributions of residual membrane potentials and distribution skews (ζ) for ongoing (top) and evoked (bottom) epochs, for traces in (a). (c) Ongoing and evoked skews for all 81 cells from 25 turtles. Each pair of connected dots indicates ongoing and evoked skews for one cell. Line above plot indicates significant change in skew from ongoing to evoked windows (P = 3.2 × 10^−4^, Wilcoxon signed-rank test). (d) Example visually-evoked high-conductance event. Red box indicates 50 ms (“pre-spike”) window ending 5 ms before spike threshold crossing (see Methods). Black box indicates 50 ms (“random”) window randomly-selected from the same high-conductance event (see Methods). (e) Pre-spike (left, top) and corresponding random (left, bottom) traces for all visually-evoked spikes for example neuron in (d). Scale bars indicate 5 mV. Each trace is fitted with a straight line via linear regression, and the slope is defined to be dV/dt. The slopes are averaged across traces for each window type, yielding >dV/dt> for pre-spike and random windows for each neuron. Right: <dV/dt> for pre-spike (red dots) and random (white dots) windows for 30 cells from 15 turtles. Asterisks above plot indicate significant difference in the populations of values (P = 1.73 × 10^−6^, Wilcoxon signed-rank test). (f) Visually-evoked spike-triggered membrane potentials for “trigger” cell (black traces) and simultaneously-recorded “paired” cell (green traces, with across-trial average in high opacity, see Methods). Short spikes are due to downsampling (see Methods). (g) Left: <dV/dt> for trigger cells (black dots) and paired cells (green dots), for 23 triggered-paired cell pairs (from 19 pairs of simultaneously-recorded cells) in 11 turtles. Note that under certain conditions, a single pair of simultaneously recorded cells can yield two trigger-paired cell pairs. Asterisks above plot indicate significant difference in the two populations of values (P = 2.70 × 10^−5^, Wilcoxon signed-rank test). Right: same as in (e, right), but for 23 paired cells from 11 turtles (P = 0.007, Wilcoxon signed-rank test).

### In the broadly-asynchronous evoked state, action potentials are preceded by concerted synaptic inputs

Having established the relatively asynchronous nature of the evoked state (at long time scales), we next asked whether the same was true at short time scales. Specifically, we asked whether synaptic inputs preceding visually-evoked spikes were consistent with an asynchronous network, in which neurons hovering just below threshold “randomly walk” the remaining distance to threshold. For each cell, we considered all spikes in a 4 s window of activity beginning 75 ms after stimulus onset (see Methods). We isolated a 50 ms (“pre-spike”) window of activity (ending 5 ms before each threshold crossing), as well as a corresponding 50 ms window of activity randomly-selected from the same 4 s window (Figure 5d). For this analysis, we required that the “random” window contain no spikes (see Methods), and that a neuron have at least six evoked spikes across all trials. In the brief pre-spike window, neurons tended to be depolarized by several millivolts (Figure 5e, **top**), in contrast to the relatively flat traces in randomly-selected windows (Figure 5e, **bottom**). Across the population, the average pre-spike depolarization far exceeded that during random windows (population grand average rate of change 
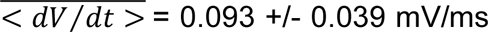
 pre-spike window, 0.002 +/− 0.014 mV/ms random window, P = 1.73 × 10^−6^ for pre-spike – random comparison, Wilcoxon signed-rank test, Figure 5e, **right**, see Methods). In other words, at short time scales (on the order of tens of milliseconds), neurons in the evoked high-conductance state were subject to coordinated excitatory synaptic inputs. This result is inconsistent with the picture of a purely asynchronous network.

### Pre-spike synaptic inputs are not strongly-correlated across nearby neurons

These pre-spike trajectories suggested coordinated spiking in presynaptic pools of neurons at short time scales. How widespread was this coordinated activity? To answer this question, we considered 19 pairs of simultaneously-recorded neurons, and inspected the subthreshold trajectories of the non-spiking (“paired”) neuron in windows preceding spikes in the “trigger” neuron (Figure 5f). In general, when one cell was driven to spike, the nearby paired neuron was subject to significantly smaller depolarization 
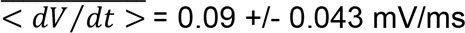
 for trigger cells, 0.01 +/− 0.02 mV/ms for paired cells, P = 2.7 × 10^−5^ for comparison, Wilcoxon signed-rank test, Figure 5f, 5g, **left**). While small, this drive to paired cells was slightly (and significantly) stronger than that during randomly-selected windows of evoked activity 
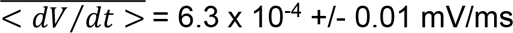
 for random windows, P = 0.007 for pre-spike – random comparison, Wilcoxon signed-rank test, Figure 5g, **right**). Thus, the synchronous, pre-spike drive to a given neuron was very nearly private to that neuron.

Taken together, these results suggest that in the evoked high-conductance state, neurons are raised toward threshold by a baseline level of synaptic input that is relatively asynchronous and non-specific across neurons at long time scales. Within this evoked state, neurons receive a final “push” to threshold by coordinated spiking in the presynaptic population. These brief windows of strong, coordinated activity are not in general common across neurons, but are instead isolated to specific microcircuits.

### Spontaneous and evoked subthreshold activity are related

We have so far considered ongoing and evoked activity separately. While all neurons were subject to large spontaneous events, the prevalence of slow-wave activity was remarkably variable across neurons (representing variability across turtles, as well as across recording sessions in a given turtle, Figure 2e; 6a, b). As this likely reflected a variability in the network state, we asked whether this attribute of spontaneous activity had any apparent impact on the visual response.

First, we quantified the prevalence of the slow-wave fluctuations in ongoing activity by calculating the fast Fourier transform (FFT) of the 9.5 s of pre-stimulus activity, and integrating over low frequencies (1 – 5 Hz), resulting in the quantity FFT_ō_ (see Methods)^18^. In agreement with qualitative inspection of voltage traces (Figure 6a, **left**; 6b, **left**), this metric was highly variable across trials (Figure 6a, **right; b, right**) and the across-trial average value (<FFT_ō_>) varied across cells (Figure 6c).

**Figure 6.**
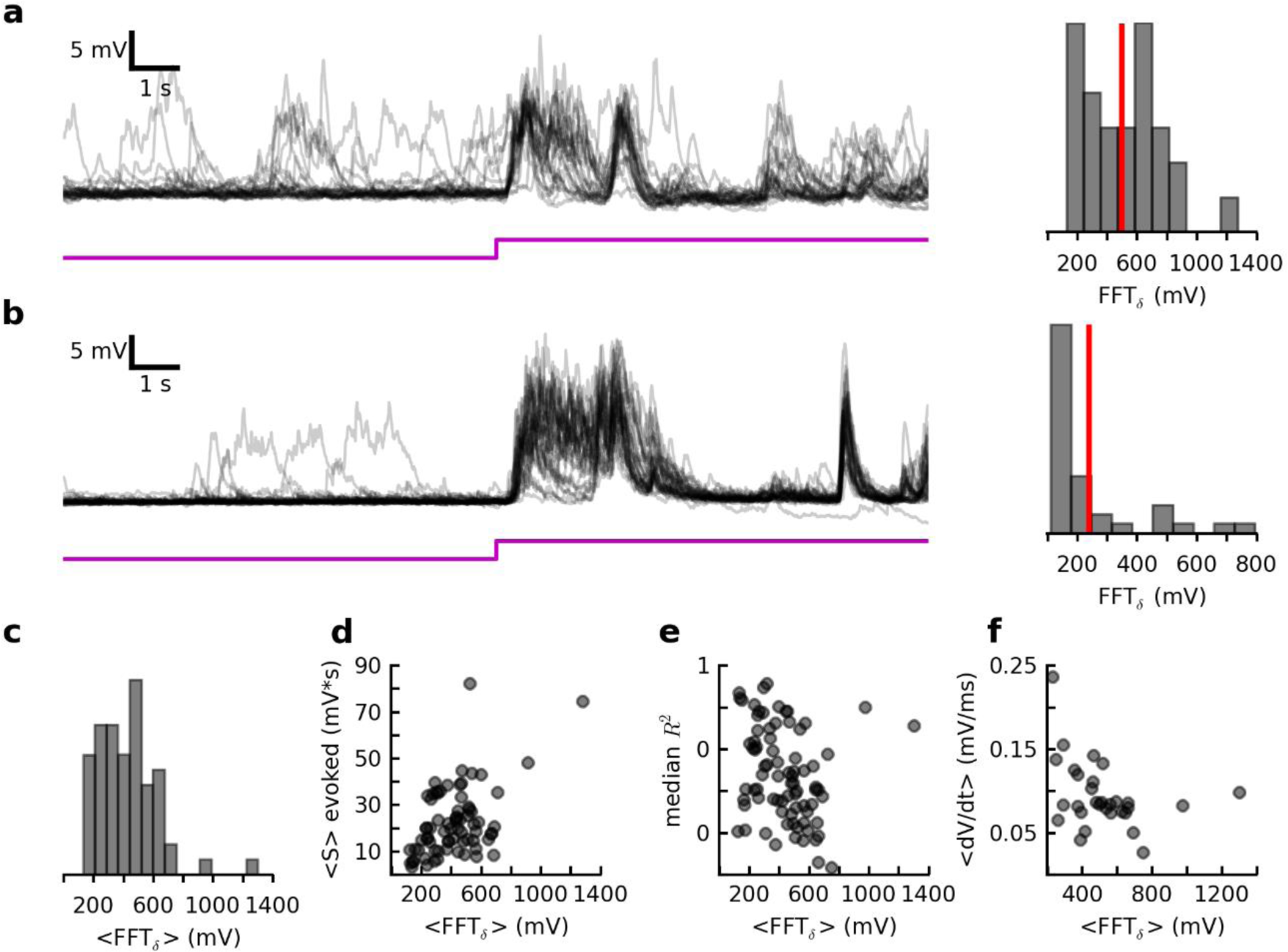
Ongoing and evoked cortical activity are related. (a – c) The prevalence of spontaneous slow-wave fluctuations varied across recording sessions and turtles. (a) Left: ongoing and evoked activity recorded from a single neuron in response to multiple presentations of a naturalistic movie (see Methods). Right: distribution of FFT_ō_ (the integrated low-frequency (1 – 5 Hz) FFT for ongoing activity) for all trials (see Methods), for cell in (a). Red vertical line indicates <FFT_ō_>, the across-trial mean. (b) Same as in (a), but for a different neuron (from the same turtle, but an earlier recording session). (c) Distribution of <FFT_ō_> values for 66 cells from 22 turtles. (d – f) The prevalence of slow-wave fluctuations in ongoing activity was related to properties of evoked activity. (d) Across-trial average subthreshold response size (<S>) vs. <FFT_ō_> for 72 cells from 23 turtles (i.e., all data points from Fig 3c, g). Ranks are significantly related (r = 0.39, P = 6.7 × 10^−4^, Spearman rank correlation). (e) Across-trial median evoked R^2^ (a proxy for response reliability, see Methods) vs. <FFT_ō_> for 79 cells from 25 turtles. Ranks are significantly related (r = -0.34, P = 1.9 × 10^−3^, Spearman rank correlation). (f) Mean evoked pre-spike depolarization (<dV/dt>) vs. <FFT_ō_> for 30 cells from 15 turtles (i.e., data points in Figure 5e, right). Ranks are not significantly related (r = -0.36, P = 0.049, Spearman rank correlation). Note: quantities are significantly linearly related when the data points corresponding to the two largest <FFT_ō_> values (i.e., the outliers in (c)) are excluded (r = -0.54, P = 0.003, linear regression). Cells used in (d – f) vary across subfigures, due to differences in requirements for calculating values on y-axis. Threshold p-value for significance has been Bonferroni-adjusted for three comparisons.

We next inspected for a relationship between the prevalence of slow-wave fluctuations (i.e.,<FFT_ō_>) and various response properties. First, we asked whether <FFT_ō_> was related to the average subthreshold response size (<S>, as in Figure 3c, g). These quantities were, in fact, positively correlated (Figure 6d). Further, these larger responses were also less reliable than smaller responses (as quantified by <R^2^>, the average variance of single-trial responses explained by the across-trial average response, Figure 6e, and see Methods). Finally, we asked whether <FFT_ō_> (indicating ongoing coordination strength at one temporal scale) could predict evoked coordination strength at shorter temporal scales (that is, the average evoked pre-spike depolarization <dV/dt>, as in Figure 6d, e). These two quantities were anti-correlated (though the coefficient was only significant when two extreme data points were removed, Figure 6f). That is, when the ongoing state was more synchronous, neurons were also closer to threshold in the evoked state, and thus required less additional depolarization to reach threshold.

Together, these results demonstrate a strong relationship between properties of the ongoing network state and those of evoked activity. Specifically, a greater prevalence of spontaneous transitions between low- and high-conductance states predicted larger and less reliable subthreshold visual responses, and smaller depolarizations immediately preceding evoked spikes.

### Visual response size depends on spontaneous activity immediately preceding the stimulus

One possible explanation for the above observations is that ongoing depolarizations add to evoked, and the random appearance of these events contributes to response variability. Alternatively, strong slow-wave activity may be consistent with a network that is more “activated” for large visual responses, yet a given spontaneous event preceding the stimulus inhibits the response.

To distinguish between these competing hypotheses, we first segregated visual stimulation trials for each cell into two categories: “low” trials, in which the stimulus was preceded by at least 2 s of quiescent ongoing activity (Figure 7a, **top**), and “high” trials, in which the stimulus either interrupted or followed soon after a large spontaneous high-conductance event (Figure 7b, **bottom**, see Methods). To be included in this analysis, we required a cell to have at least three “high” trials. This effectively restricted the set of included cells to those displaying relatively prominent slow-wave ongoing activity. We then calculated the subthreshold response size scaled by the maximum across-trial response size (S’/S’_max_) in a 2 s post-stimulus window, and averaged across trials (see Methods). We found that “high” responses were significantly smaller than “low” responses (population grand average response amplitude 
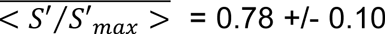
 for “low” responses, 0.68 +/− 0.14 for “high” responses, P = 4.52 × 10^−4^ for comparison, Wilcoxon signed-rank test, Figure 7b). Evidently, while a cortex with more frequent spontaneous high-conductance events yielded larger average responses (Figure 6d), a given visual response was, in fact, inhibited by excessive activity in the window immediately preceding the stimulus.

**Figure 7.**
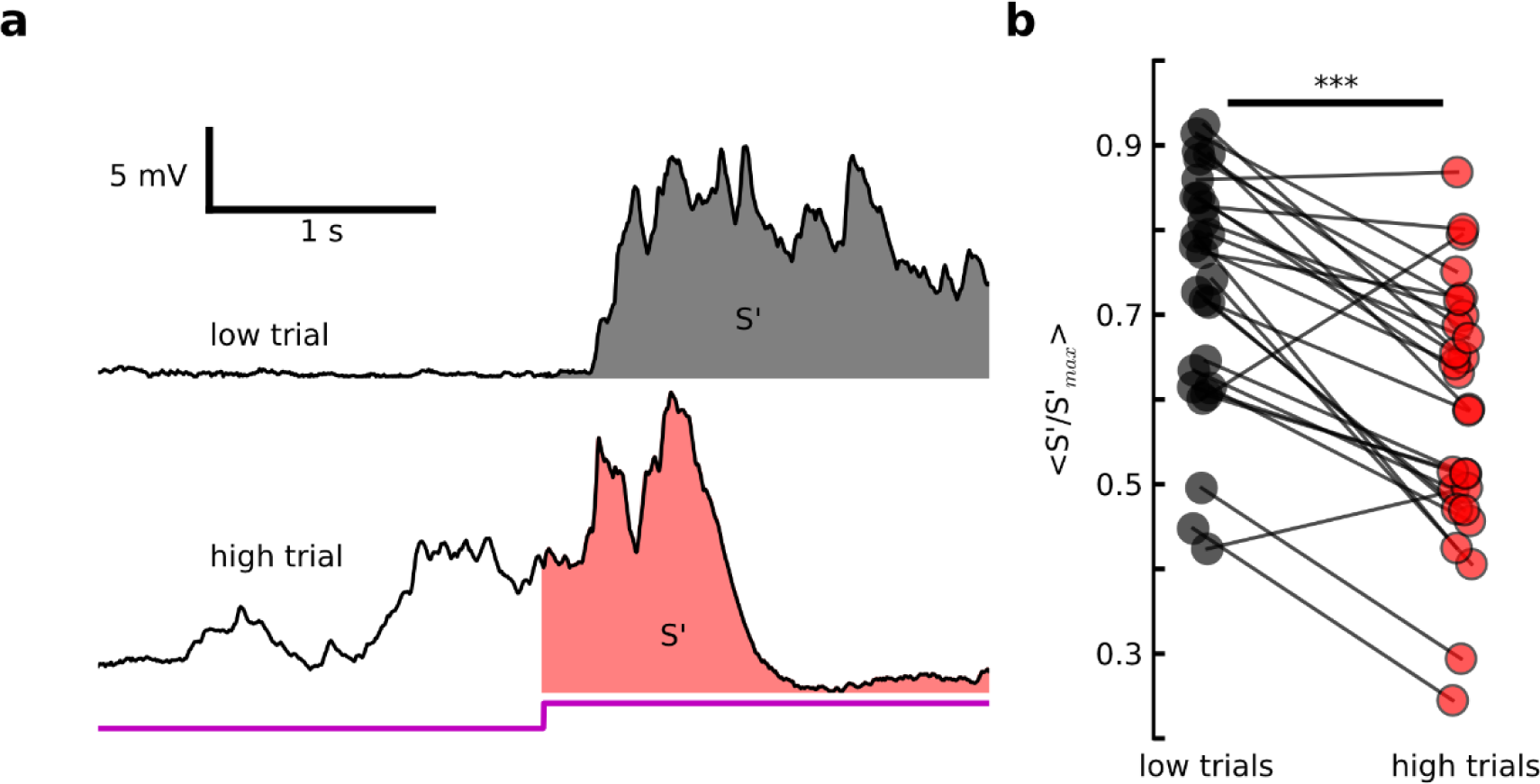
Large, spontaneous pre-stimulus events correspond to smaller visual responses. (a) Two example responses from one cell to presentation of naturalistic movie (see Methods). Top: “low trial”, in which stimulus presentation is preceded by low-conductance state. Bottom: “high trial”, in which stimulus is preceded by or interrupts a large, spontaneous high-conductance event. S’ is the area under the curve for the 2 s after stimulus onset. (b) Average scaled S’ (see Methods) for low (black dots) and high (red dots) trials for 27 cells from 15 turtles. Asterisks above plot indicate significant difference between the two populations of values (<S’/S’max> = 0.73 +/− 0.14 for low trials, 0.58 +/− 0.15 for high trials, P = 6.01 × 10^−5^ for low-high comparison, Wilcoxon signed-rank test.

## Discussion

In order to investigate ongoing and visually-evoked subthreshold and spiking cortical activity, we obtained whole-cell recordings from neurons in the three-layer visual cortex of the *ex vivo* turtle eye-attached whole-brain preparation. To infer properties of the corresponding network activity, we investigated the statistics of subthreshold activity at two distinct time scales for individual neurons, and for pairs of simultaneously-recorded neurons, and compared to results predicted by theory. Finally, we investigated the relationship between ongoing and evoked network activity by comparing properties of the two network states (as communicated by subthreshold activity).

In the absence of visual stimulation, neurons were usually in a relatively quiescent “low-conductance” state, with random transitions into depolarized “high-conductance” states (Figure 2a). These are often referred to as “Down” and “Up” states, respectively, but the conductance-based labels underscore the synaptic origin of these events, which signal changes in the activity level of a neuron’s presynaptic pool. While drastic Up-Down transitions can be brought on by anesthesia^19^, many studies have demonstrated similar non-Gaussian spontaneous fluctuations in a variety of areas in awake preparations, including mouse^12^ and primate^4^ visual cortex, rat auditory cortex^3^, rodent somatosensory cortex^13–16^, and even in reduced preparations, including thalamocortical slice^20–23^. Previous work has shown that the transient increases in presynaptic firing are due to waves of activity propagating across the cortex^15^. These waves (also observed in turtle cortex^24^) are thought to be generated intracortically^23,25^, though they can be triggered and shaped by external inputs^21,26–28^. Recent computational work suggests that the cortical excitation/inhibition balance is crucial to this phenomenon^29^. Thus, the spontaneous membrane potential dynamics we observe here likely reflect the complex spatiotemporal population activity patterns that emerge in balanced cortical networks.

Visual stimulation evoked high-conductance states (Figure 3a, e) that were qualitatively similar to longer-duration spontaneous high-conductance events (Figure 2a,b, d). It is possible there is more than a passing resemblance between these two types of activity. Previous work has uncovered striking similarities between ongoing and evoked activity in cortex^30–32^, and specifically in the subthreshold activity of individual cortical neurons^21^. Because a neuron’s subthreshold time series communicates a spatiotemporal sequence of cortical firing, this indicates similar spontaneous and evoked firing patterns. Such observations are consistent with the view that spontaneous cortical events broadly sample the set of all possible spatiotemporal patterns in the cortical “response manifold”^33^. While we did not perform an in-depth investigation of this topic, this preparation has great potential for such a study. First, it represents an important advance beyond the slice preparation; intracortical connectivity is preserved, and cortical activity is evoked by visual (rather than electrical) stimulation. Second, multi-whole-cell recordings allow for a “higher-dimensional” view of cortical firing than that afforded by individual whole-cell recordings. Future work can therefore enhance our understanding of the relationship between ongoing and evoked cortical activity.

Previous studies have described a stimulus-induced transition from cortical synchrony to asynchrony^4^. In other words, pre-stimulus subthreshold activity includes frequent periods of broadly-correlated depolarization, and evoked activity more closely follows the “random walk” dynamics consistent with Poisson process inputs. Yet in other studies, sensory-evoked spikes are preceded by strong depolarizations, indicating concerted presynaptic firing^14^,^16^. We observed both of these seemingly-contradictory phenomena (Figure 5a-e). This paradox is actually predicted for neurons in the high-conductance state; while the inputs to a neuron may be relatively random, the reduced membrane time constant make the neuron a better coincidence detector^19,34^. Thus, evoked spikes in a given neuron are more likely to result from nearly-coincident synaptic inputs, and to be more tightly phase-locked to those of the presynaptic population. This mechanism for spike synchronization is most effective when a neuron’s reduced time constant (promoting coincidence detection) outweighs its reduced distance from spike threshold (which makes the depolarized cell more likely to spike in response to even small additional inputs). We found that the balance between these two biophysical influences depended on cortical state; pre-spike depolarization was smaller when the ongoing cortical state was consistent with larger depolarizations (Figure 6d, f), suggesting the coincidence-detection mechanism was more easily “saturated”. Our observations thus provide a rare, intracellular view of the relationship between long-time-scale coordination during ongoing activity and short-time-scale coordination (leading to spikes) during visual processing.

Pairwise recordings revealed a spatial dimension to coordination as well; at a fixed temporal scale (tens of milliseconds), activity could become strongly coordinated in a given microcircuit (Figure 5d), but was not, in general, coordinated across cortical space (Figure 5f, g). This is likely due at least in part to the nature of cortical connectivity; dense interconnectivity supports broadly-distributed states of enhanced excitability (that is, broadly-coordinated transitions to high-conductance states), while the “constellation-like” specificity of this connectivity (with a minority of extremely strong connections, and a majority of weak ones^35^) diversifies the coupling strengths among neurons in a local population^36,37^. This may create preferred pathways for the propagation of spikes within the high-conductance state. Combined with the observed temporal dependence, this result demonstrates just two of the possibly many spatiotemporal scales at work in the likely “multiplexed” cortical code^38^.

While anatomical connectivity is extremely influential to evoked activity, the ongoing network state is also thought to play an important role. For instance, some set of conditions hidden from the experimenter modulates the degree of ongoing cortical slow-wave activity (or synchrony). How do the variables controlling these properties of ongoing activity affect sensory responses? Previous studies have reported a rich dependence of cortical response properties on the spontaneous cortical state^18,39–41^. Accordingly, we found that stronger slow-wave fluctuations predicted visual responses that were on average larger (Figure 6d) and less reliable (Figure 6e) than those corresponding to more quiescent ongoing states. While we did not directly investigate the deeper implications, two results suggest a nonlinear relationship between the ongoing state and cortical function; the enhanced variability at the high end of the synchrony “spectrum” seems disadvantageous for sensory encoding, while the smaller responses at the other end may be as well. The optimal state may be an intermediate level of synchrony, balancing these two phenomena. In support of this idea, recent work has suggested that maximum dynamic range occurs in a cortex tuned to the critical state^42^, which also coincides with intermediate levels of network synchrony^43,44^. Where the cortex sits on this spectrum is likely governed by top-down control, most generally tracked by level of arousal. Indeed, intermediate levels of arousal correspond to intermediate spontaneous membrane potential levels and enhanced perception^45^. It will be important for future work to identify such general principles of cortical function (e.g., criticality) that govern phenomena commonly observed in experiment (e.g., synchrony, as observed in the statistics of population spiking or subthreshold membrane potential fluctuations), as well as the synaptic basis of top-down interactions.

Finally, there is the question of how the condition of the cortex at or immediately preceding stimulus onset influences the response. This question becomes increasingly relevant as the ongoing network state becomes more synchronous, as large, spontaneous events become more common. Many studies have observed that the probabilistic nature of ongoing activity contributes to across-trial response variability^19,26,41,46^. The exact nature of the interaction is an item of debate^47^. On the one hand, spontaneous depolarization brings neurons closer to threshold prior to the arrival of excitatory sensory input. Accordingly, some studies show that enhanced levels of pre-stimulus activity correspond to larger responses^46,48–50^. On the other hand, spontaneous synaptic barrages reduce input resistance^19^,^51,52^ and depress synapses^53–55^. In addition, spontaneous depolarization increases the driving force for inhibition^55^, and thus the IPSP amplitudes associated with short-latency, disynaptic feedforward inhibiton^56^. Consistent with this view, some studies have reported that large spontaneous events suppress evoked activity^15,57^. Our results are in agreement with these latter studies; when the visual stimulus interrupted or followed soon after a large spontaneous event, the subthreshold response was muted (Figure 7). Evidently, suppressing mechanisms (likely including those described above) more than counterbalanced the reduced distance to threshold in the presynaptic population.

What are the implications of this dependency on the pre-stimulus condition? Previous work provides conflicting answers. In some cases, a high-conductance pre-stimulus state corresponds to muted responses that are less reliable across trials^13^, thus compromising response fidelity. Still, in barrel cortex, this muted responses (resulting from either spontaneous pre-stimulus events^15,58^, or the presentation of a background stimulus^59,60^) are more confined to the column corresponding to the stimulated whisker, which promotes stimulus-response mutual information. Elsewhere, a more active pre-stimulus condition yields larger responses, that are less effective at transmitting information^50^. Finally, at least one study shows no measurable relationship between the pre-stimulus state and the size or reliability of the early cortical response^18^. Of course, another possibility is that the “adapted” state resulting from prominent pre-stimulus activity is optimized for some functions (e.g., discrimination) at the expense of others (e.g., detection)^59,60^. Evidently, it will be important for future studies to address the impact of the pre-stimulus state on sensory responses using carefully-designed stimuli and information-theoretic measures of both detection and discrimination. As in the discussion of long-time-scale ongoing “state”, it will be important to incorporate behavior; a recent study has shown that diminished “late” responses (corresponding to stimuli delivered in the Up state) causally impair perception^18^. This combined approach can be extremely challenging to implement (especially when involving patch clamp recording), but is becoming increasingly feasible. While such techniques are being developed, it will be important to continue to document the effects of the pre-stimulus cortical state on sensory responses to the extent possible; this will reveal which aspects of the interaction generalize across areas and species.

In conclusion, these results contribute to a clearer picture of the subthreshold dynamics of cortical visual responses. They highlight the importance of the relationship between the ongoing network state and subthreshold evoked activity. Further, they show that evoked spiking is shaped by presynaptic activity that is coordinated at multiple spatiotemporal scales. As such, this study is in agreement with previous work suggesting that anatomical and emergent cortical network properties play vital roles in cortical sensory processing, and provides a rare view of this influence at the level of the membrane potential.

## Methods

### Surgery

All procedures were approved by Washington University’s Institutional Animal Care and Use Committees and conform to the guidelines of the National Institutes of Health on the Care and Use of Laboratory Animals. Fourteen adult red-eared sliders (*Trachemys scripta elegans*, 150-1000 g) were used for this study. Turtles were anesthetized with Propofol (2mg Propofol/kg), then decapitated. Dissection proceeded as described previously^7,8,61^. In brief, immediately after decapitation, the brain was excised from the skull, with right eye intact, and bathed in cold extracellular saline (in mM, 85 NaCl, 2 KCl, 2 MgCl_2_*6H_2_O, 20 Dextrose, 3 CaCl_2_-2H_2_O, 45 NaHCO_3_). The dura was removed from the left cortex and right optic nerve, and the right eye hemisected to expose the retina. The rostral tip of the olfactory bulb was removed, exposing the ventricle that spans the olfactory bulb and cortex. A cut was made along the midline from the rostral end of the remaining olfactory bulb to the caudal end of the cortex. The preparation was then transferred to a perfusing chamber (Warner RC-27LD recording chamber mounted to PM-7D platform), and placed directly on a glass coverslip surrounded by Sylgard. A final cut was made to the cortex (orthogonal to the previous and stopping short of the border between medial and lateral cortex) allowing the cortex to be pinned flat, with ventricular surface exposed. Multiple perfusion lines delivered extracellular saline, adjusted to pH 7.4 at room temperature, to the brain and retina in the recording chamber.

### Intracellular Recordings

We performed whole-cell current clamp recordings from 39 cells in 14 preparations. Patch pipettes (4-8 MΩ) were pulled from borosilicate glass and filled with a standard electrode solution (in mM; 124 KMeSO_4_, 2.3 CaCl_2_-2H_2_O, 1.2 MgCl_2_, 10 HEPES, 5 EGTA) adjusted to pH 7.4 at room temperature. Cells were targeted for patching using a dual interference contrast microscope (Olympus). All cells were located within 300 microns of an extracellular recording electrode. Intracellular activity was collected using an Axoclamp 900A amplifier, digitized by a data acquisition panel (National Instruments PCIe-6321), and recorded using a custom Labview program (National Instruments), sampling at 10 kHz. The visual cortex was targeted as described previously^61^.

### Extracellular Recordings

We performed extracellular recordings at 12 recording sites in seven preparations. We used tungsten microelectrodes (MicroProbes heat treated tapered tip), with approximately 0.5 MΩ impedance. Electrodes were slowly advanced through tissue under visual guidance using a manipulator (Narishige), while monitoring for spiking activity using custom acquisition software (National Instruments). Extracellular activity was collected using an A-M Systems Model 1800 amplifier, band-pass filtered between 1 Hz and 20,000 Hz, digitized (NI PCIe-6231), and recorded using custom software (National Instruments), sampling at 10 kHz.

### Visual Stimulation

Whole-field flashes were presented using either a red LED (Kingbright, 640nm), mounted to a manipulator and positioned 1 – 5 cm above the retina, or a projector-lens system (described below). The mean LED light intensity (irradiance) at the retina was 60 W/m^2^. For one turtle, we used these same LEDs in conjunction with 200 micron optical fibers (Edmund Optics) to project sub-field flashes (1 ms – 200 ms) onto the visual streak. Other stimuli were presented using using a projector (Aaxa Technologies, P4X Pico Projector), combined with a system of lenses (Edmund Optics) to project images generated by a custom software package directly onto the retina. The mean irradiance at the retina was 1 W/m^2^. This system was used to present brief (100 ms – 250 ms) whole-field and sub-field flashes (red or white), sustained (10 s) gray screen, a naturalistic movie (“catcam”) a motion-enhanced movie (courtesy Jack Gallant), and a phase-shuffled version of the same movie (courtesy Jack Gallant and Woodrow Shew). In all cases, the stimulus was triggered using a custom Labview program (National Instruments).

For each cell and extracellular recording site, we selected one of the five stimuli listed above to present across all trials. The preparation was in complete darkness before and after each stimulus presentation. Extended stimuli lasted either 10 s or 20 s, and flashes lasted between 1 ms and 250 ms, with at least 30 s between the end of one presentation and the beginning of the next. In all cases, visual stimulation trials were repeated at least 12 times.

### Data included in analysis

For each extracellular recording site, we used visual inspection to determine the quality of the recordings. In general, we excluded recording sites from consideration if voltage traces displayed excessive 60 Hz line noise, low-frequency noise (likely reflecting a damaged electrode), or on average small response amplitudes relative to baseline.

For the analysis of intracellular recordings, we required at least twelve visual stimulation trials (unless stated otherwise, see below).

### Processing of intracellular and extracellular voltage recordings

Raw data traces were down-sampled to 1000 Hz. We used an algorithm to detect spikes in the membrane potential, and the values in a 20 ms window centered on the maximum of each spike were replaced via interpolation. Finally, we applied a 100 Hz lowpass Butterworth filter. We did not perform these last two steps for “trigger cells” used to calculate spike-triggered averages (Figure 5d-g, and see below)

In addition, we detrended spontaneous recordings. To do this, for each time step, we subtracted a value obtained from a 10 s window beginning at that time step. The value we used depended on the median membrane potential (MMP) of the cell in each full recording; if the MMP was above −60 mV, we used the median value of the 10 s window for detrending. If MMP was below −60 mV, we used the fifth percentile of the window. The reasoning for this was as follows: in the detrending process, we seek to subtract from the membrane potential at a given point in time a good estimate of the “true” resting membrane potential (RMP) near that point in time. When MMP is low, high-conductance events usually result in large depolarizations. Even when using large sliding windows in the detrending process, these depolarizations can lead to spurious changes in the detrended membrane potential before and after each event. We therefore use the fifth percentile of the window for low-MMP cells, as it is less susceptible than the median to outlier membrane potential values above MMP, and is therefore a better estimate of RMP in the neighborhood of the event. For neurons with high MMP, on the other hand, IPSPs can be quite large during high-conductance events (due to the increased distance from the inhibitory reversal potential). In fact, the membrane potential often drops below MMP during these events, meaning the fifth percentile does as well. In this case, the median of the window is a better estimate of RMP in the neighborhood of the event.

### Detecting and quantifying spontaneous high-conductance events

We used an algorithm to detect spontaneous high-conductance events in each detrended spontaneous voltage trace (Figure 5b, **top**). First, we detected all “bumps” (i.e., windows of activity in which the membrane potential exceeded the standard deviation of the full trace by at least a factor of 1.5). Then, we detected “high bumps” (using instead a threshold factor of 4). Finally, given these “bumps” and “high bumps”, we identified “high-conductance events”. First, we combined “bumps” separated by less than 500 ms of silence. Any “bump” in this refined set that also included a “high bump” was defined to be a “high-conductance event”. For each event (of duration D in ms), we defined the size (S in mV*s) to be the area under the curve (Figure 5b, **bottom**).

### Subthreshold response latencies

For each presentation of a stimulus (flashes and extended stimuli), we calculated response latency (Figure 3b, f) by first considering the membrane potential in a window of activity beginning 9 s before and ending 9 s after stimulus onset. We then calculated the slope (dV/dt) as a function of time for this sub-trace. We excluded the trial from consideration if 1) visual stimulation clearly interrupted a large spontaneous Up state, or 2) large PSPs were present immediately before stimulus onset (indicating the possible onset of a spontaneous high-conductance event). We considered condition 1 (2) to be satisfied if any value of the slope trace in the 500 ms (200 ms) preceding stimulus onset exceeded six (four) times the standard deviation of the entire slope trace. If neither of these conditions were met, we defined the response latency to be the time after stimulus onset at which the slope trace exceeded three times the standard deviation of the entire slope trace. Finally, “latencies” smaller than 50 ms were excluded.

### Subthreshold response duration

To calculate the duration (D) of a subthreshold flash response (Figure 3c), we first applied the high-conductance event detection algorithm (described above for spontaneous events) to a window (18 s centered on the stimulus onset) extracted from the full voltage trace. With the resulting set of event times (i.e., event onset, offset pairs), we considered the response onset to be the first event onset time in the window between 50 ms and 1 s after stimulus onset. The response offset was the last event offset time in a window between response onset and 7 s after stimulus onset. Only trials with valid onset and offset times were included.

### Subthreshold response size

To calculate subthreshold response size (S) for flashes (Figure 3d) and extended stimuli (Figure 3g), we first detected the response duration (as described above). Then, for each response window, we subtracted the fifth percentile of the membrane potential (calculated from the full trial). We defined the response size to be the area under the curve for the response window. Note that for extended stimuli, activity typically persisted beyond the window used to calculate response size. For this type of stimulus, then, S is a measure of evoked activity in at most the first 7 s of the response (and is still a useful measure for comparisons across events (Figure 3g, **left**) and cells (Figure 3g, **right**) for extended stimuli only).

### Evoked action potential rates

To identify spike times in intracellular recordings of visual responses (Figure 4a), we first estimated the first derivative of the voltage trace (V’) by calculating the change in membrane potential for each (1 ms) time step. We then defined spike times to be those at which the value of V’ was at least 20 times the standard deviation of V’. Finally, we calculated the average spike rate for each cell in a two-second window after stimulus onset (Figure 4a, **inset**), which was a window that contained most of the evoked spikes in a typical trial (Figure 4b, c).

### Multi-unit activity

We determined multi-unit spiking activity for one recording session, to provide an example of spiking patterns in a population nearby two intracellularly-recorded neurons (Figure 4a, **bottom**). Multi-unit spike times for each trial were defined to be those at which the value of the high-pass-filtered (250 Hz Butterworth) extracellular trace was at least six times the standard deviation of the full trace.

### Peristimulus time histograms

We constructed peristimulus time histrograms (PSTHs) for responses to brief (Figure 4b) and extended (Figure 4c) stimuli. For each stimulus, we pooled the responses from all intracellular recordings. For each trial, spike times were detected as described above. We included all trials that contained at least one spike in the 7 s after stimulus onset.

### Residual membrane potentials and residual skew

For each recorded cell, we calculated the residual time series (single-trial time series with across-trial average time series subtracted, Figure 5a). We then considered two windows of activity: the ongoing (2 s to 0 s before stimulus onset), and evoked (0 s to 2s after response onset) windows. Response onset was determined as described above (for response latency calculations). For each trial and epoch, we subtracted the average value for that epoch, and concatenated the resulting trace to a single time series. We then calculated the skew of the concatenated time series (Figure 5b). We compared the results for all cells (Figure 5c) using the Wilcoxon signed-rank test.

### Spike-triggered membrane potentials

For each cell and visual stimulation trial, we determined evoked action potential times, as described above. An action potential was included in this analysis if it occurred between 75 ms and 4 s after stimulus onset. We included a cell in this analysis if it spiked at least six times in this response window across all trials. For each spike, the threshold crossing time was defined to be the first zero-crossing of the second time derivative of V in the 5 ms preceding the spike time. This represents the maximum change in membrane potential slope immediately before the spike. We estimated the second derivative (V’’) at each time step *k* using Taylor series expansion^62^:

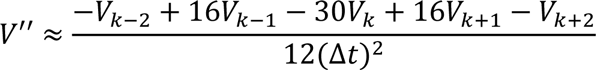

where Δ*t* is the size of the time step (1 ms).

We then considered a 50 ms “pre-spike” window of activity ending 5 ms before threshold crossing. For each spike, we also considered a 50 ms “random” window of activity that (i) was taken from the same 4 s response window; and (ii) did not contain an action potential (Figure 5d). We found the slope of the trace in each window via linear regression, and then averaged across trials (yielding <dV/dt> for pre-spike and random windows, Figure 5e). We compared <dV/dt> for the two window types using the Wilcoxon signed-rank test.

We performed a similar analysis for pairs of simultaneously-recorded neurons. For each pair, if either of the cells had at least six evoked spikes across all trials, it was treated as the “trigger” cell, and the other as the “paired” cell (Figure 6f). Average “pre-spike” and “random” depolarizations (<dV/dt>) were calculated as described above, using the spike times from the trigger cell. We compared <dV/dt> values for trigger and paired cells (Figure 6g, **left**), and for pre-spike and random windows for paired cells (Figure 6g, **right**) using the Wilcoxon signed-rank test.

### Low-frequency FFT of ongoing activity

For each cell, we considered the ongoing activity in the 9.5 s before stimulus onset (Figure 6a, b), and calculated the fast Fourier transform (implemented using the numpy.fft routine, with documentation available at <u>https://docs.scipy.org/doc/numpy/reference/generated/numpy.fft.fft.html</u>). We averaged the sum of the 1 – 5 Hz Fourier coefficients across all trials, yielding <FFTō>, Figure 6c), which, for a given cell, was a reliable proxy for the prevalence of large, spontaneous high-conductance events in the pre-stimulus window (Figure 6a, b).

### Response reliability

For each cell, we considered a 2 s window of evoked activity beginning at stimulus onset. For this window, we regressed each single-trial response onto the average response, yielding R^2^ (the explained variability). For each cell, we took the across-trial median R^2^ value (Figure 6e) to be the “response reliability”.

### “Low” and “high” visual response trials and scaled response size

We segregated the visual responses for each cell into two categories defined by ongoing activity: “low” and “high” trials. For each recording, we used an algorithm to identify all high-conductance events (as described above). The trial was designated a “high” trial if a spontaneous high-conductance event (i) ended within 2 s of stimulus onset; or (ii) was interrupted by stimulus onset (Figure 7a). A cell was required to have at least three high and three low trials to be included in this analysis. We forced the two trial sets to contain the same number of trials by randomly excluding trials from the longer of the two sets. Of these remaining trials, we considered a 2s window of activity beginning at stimulus onset. From this evoked window, we subtracted a baseline value (the fifth percentile, calculated from the full, original voltage trace). We defined the response size (S’) to be the area under the resulting curve (Figure 7a). We scaled each response size by the maximum response size (S’max) in the retained trials, and averaged across trials (yielding <S’/S’max>. We then compared the populations of average scaled responses using the Wilcoxon signed-rank test (Figure 7b). Because the process of excluding trials was random, we ensured the significant difference indicated in Figure 7b was robust to iterations of the calculation (data not shown).

### Statistical analysis

All statistical tests were performed using Python 2.7.

Before applying any significance test that assumed normality, we performed an omnibus test for normality on the associated dataset(s). This test compares the skew and kurtosis of the population from which the dataset was drawn to that of a normal distribution, returning a p-value for a two-sided chi-squared test of the null hypothesis that the data is drawn from a normal distribution. This test is valid for sample sizes of 20 or larger, and was implemented using scipy.stats.mstats.normaltest (documentation and references available at http://docs.scipy.org/doc/scipy-0.14.0/reference/generated/scipy.stats.mstats.normaltest.html). We report these p-values as the result of a “two-sided omnibus chi-shared test for normality”.

When asking whether a parameter of interest changed significantly across two sets of conditions for a population, we applied the Wilcoxon signed-rank test, which returns a p-value for the two-sided test that the two related paired samples are drawn from the same distribution. This test assumes normality, and was implemented using scipy.stats.wilcoxon (documentation and references available at <u>http://docs.scipy.org/doc/scipy/reference/generated/scipy.stats.wilcoxon.html</u>).

## Acknowledgments

We thank Woodrow Shew for assistance with the design of the visual stimulus. We thank Thomas Crockett for contributing data. This research was supported by a Whitehall Foundation grant #20121221 (R.W.) and a NSF CRCNS grant #1308159 (R.W.).

## Author Contributions

N.W. and R.W. conceived the study and designed the experiments. N.W. performed the experiments and analyzed the data. N.W. and R.W. wrote the paper.

## Competing Financial Interests

The authors declare no competing financial interests.

## References

1. Shoham, S., O’Connor, D. H. & Segev, R. How silent is the brain: Is there a ‘dark matter’ problem in neuroscience? J. Comp. Physiol. A Neuroethol. Sensory, Neural, Behav. Physiol. 192, 777–784 (2006).

2. Cohen, M. R. & Kohn, A. Measuring and interpreting neuronal correlations. Nat. Neurosci. 14, 811–819 (2011).

3. DeWeese, M. R. & Zador, A. M. Non-Gaussian Membrane Potential Dynamics Imply Sparse, Synchronous Activity in Auditory Cortex. J. Neurosci. 26, 12206–12218 (2006).

4. Tan, A. Y. Y., Chen, Y. Scholl, B., Seidemann, E. & Priebe, N. J. Sensory stimulation shifts visual cortex from synchronous to asynchronous states. Nature 509, 226–229 (2014).

5. Rudolph, M., & Destexhe, A. Characterization of subthreshold voltage fluctuations in neuronal membranes. Neural Comput. 15, 2577–618 (2003).

6. Stevens, C. F. & Zador, a M. Input synchrony and the irregular firing of cortical neurons. Nat. Neurosci. 1, 210–7 (1998).

7. Saha, D., Morton, D. Ariel, M. & Wessel, R. Response properties of visual neurons in the turtle nucleus isthmi. J. Comp. Physiol. A 197, 153–165 (2011).

8. Crockett, T., Wright, N. Thornquist, S., Ariel, M. & Wessel, R. Turtle Dorsal Cortex Pyramidal Neurons Comprise Two Distinct Cell Types with Indistinguishable Visual Responses. PLoS One 10, e0144012 (2015).

9. Fournier, J., Müller, C. M. & Laurent, G. Looking for the roots of cortical sensory computation in three-layered cortices. Curr. Opin. Neurobiol. 31, 119–126 (2015).

10. Naumann, R. K. et al. The reptilian brain. Curr. Biol. 25, R317–R321 (2015).

11. Shepherd, G. M. The microcircuit concept applied to cortical evolution: from three layer to six-layer cortex. Front. Neuroanat. 5, 30 (2011).

12. Bennett, C., Arroyo, S. & Hestrin, S. Subthreshold mechanisms underlying state dependent modulation of visual responses. Neuron 80, 350–7 (2013).

13. Crochet, S., & Petersen, C. C. H. Correlating whisker behavior with membrane potential in barrel cortex of awake mice. Nat. Neurosci. 9, 608–610 (2006).

14. Gentet, L. J., Avermann, M. Matyas, F., Staiger, J. F. & Petersen, C. C. H. Membrane potential dynamics of GABAergic neurons in the barrel cortex of behaving mice. Neuron 65, 422–35 (2010).

15. Petersen, C. C. H., Hahn, T. T. G., Mehta, M. Grinvald, A. & Sakmann, B. Interaction of sensory responses with spontaneous depolarization in layer 2/3 barrel cortex. Proc. Natl. Acad. Sci. U. S. A. 100, 13638–43 (2003).

16. Poulet, J. F. a & Petersen, C. C. H. Internal brain state regulates membrane potential synchrony in barrel cortex of behaving mice. Nature 454, 881–5 (2008).

17. Marchiafava, P. L. An “antagonistic” surround facilitates central responses by retinal ganglion cells. Vision Res. 23, 1097–1099 (1983).

18. Sachidhanandam, S., Sreenivasan, V. Kyriakatos, A., Kremer, Y. & Petersen, C. C. H. Membrane potential correlates of sensory perception in mouse barrel cortex. Nat. Neurosci. 16, 1671–7 (2013).

19. Destexhe, A., Rudolph, M. & Paré, D. The high-conductance state of neocortical neurons in vivo. Nat. Rev. Neurosci. 4, 739–751 (2003).

20. Cossart, R., Aronov, D. & Yuste, R. Attractor dynamics of network UP states in the neocortex. 423, 13–16 (2003).

21. MacLean, J. N., Watson, B. O. Aaron, G. B. & Yuste, R. Internal dynamics determine the cortical response to thalamic stimulation. Neuron 48, 811–823 (2005).

22. Graupner, M., & Reyes, A. D. Synaptic Input Correlations Leading to Membrane Potential Decorrelation of Spontaneous Activity in Cortex. J. Neurosci. 33, 15075–15085 (2013).

23. Sanchez-Vives, M. V & McCormick, D. A. Cellular and network mechanisms of rhythmic recurrent activity in neocortex. Nat. Neurosci. 3, 1027–1034 (2000).

24. Senseman, D. M. & Robbins, K. a. Modal behavior of cortical neural networks during visual processing. J. Neurosci. 19, RC3 (1999).

25. Steriade, M., Contreras, D. Curró Dossi, R. & Nuñez, A. The slow (< 1 Hz) oscillation in reticular thalamic and thalamocortical neurons: scenario of sleep rhythm generation in interacting thalamic and neocortical networks. J. Neurosci. 13, 3284–99 (1993).

26. Hirata, A., & Castro-Alamancos, M. a. Effects of cortical activation on sensory responses in barrel cortex. J. Neurophysiol. 105, 1495–1505 (2011).

27. Rigas, P., & Castro-Alamancos, M. A. Thalamocortical Up states: differential effects of intrinsic and extrinsic cortical inputs on persistent activity. J. Neurosci. 27, 4261–72 (2007).

28. Poulet, J. F. A., Fernandez, L. M. J., Crochet, S. & Petersen, C. C. H. Thalamic control of cortical states. Nat. Neurosci. 15, 370–372 (2012).

29. Keane, A., & Gong, P. Propagating waves can explain irregular neural dynamics. J Neurosci 35, 1591–1605 (2015).

30. Sakata, S., & Harris, K. D. Article Laminar Structure of Spontaneous and Sensory Evoked Population Activity in Auditory Cortex. Neuron 64, 404–418 (2009).

31. Kenet, T., Bibitchkov, D. Tsodyks, M., Grinvald, A. & Arieli, A. Spontaneously emerging cortical representations of visual attributes. Nature 425, 954–956 (2003).

32. Luczak, A., Barthó, P. & Harris, K. D. Spontaneous Events Outline the Realm of Possible Sensory Responses in Neocortical Populations. Neuron 62, 413–425 (2009).

33. Ringach, D. L. Spontaneous and driven cortical activity: implications for computation. Curr. Opin. Neurobiol. 19, 439–44 (2009).

34. Rudolph, M., & Destexhe, A. A Fast-Conducting , Stochastic Integrative Mode for Neocortical Neurons In Vivo. 23, 2466–2476 (2003).

35. Cossell, L., et al. Functional organization of excitatory synaptic strength in primary visual cortex. Nature 0, 1–5 (2015).

36. Okun, M., et al. Diverse coupling of neurons to populations in sensory cortex. Nature 521, 511–515 (2015).

37. Hofer, S. B. et al. Differential connectivity and response dynamics of excitatory and inhibitory neurons in visual cortex. Nat. Neurosci. 14, 1045–52 (2011).

38. Panzeri, S., Brunel, N. Logothetis, N. K. & Kayser, C. Sensory neural codes using multiplexed temporal scales. Trends Neurosci. 33, 111–120 (2010).

39. Ecker, A. S. et al. State dependence of noise correlations in macaque primary visual cortex. Neuron 82, 235–48 (2014).

40. Haider, B., Schulz, D. P. A. P. A., Carandini, M. Häusser, M. & Carandini, M. Millisecond Coupling of Local Field Potentials to Synaptic Currents in the Awake Visual Cortex. Neuron 90, 35–42 (2015).

41. Scholvinck, M. L., Saleem, A. B. Benucci, A., Harris, K. D. & Carandini, M. Cortical State Determines Global Variability and Correlations in Visual Cortex. J. Neurosci. 35, 170–178 (2015).

42. Shew, W. L., Yang, H. Petermann, T., Roy, R. & Plenz, D. Neuronal Avalanches Imply Maximum Dynamic Range in Cortical Networks at Criticality. J. Neurosci. 29, 15595–15600 (2009).

43. Gautam, S. H., Hoang, T. T. McClanahan, K., Grady, S. K. & Shew, W. L. Maximizing Sensory Dynamic Range by Tuning the Cortical State to Criticality. PLoS Comput. Biol. 11, 1–15 (2015).

44. Yang, H., Shew, W. L. Roy, R. & Plenz, D. Maximal Variability of Phase Synchrony in Cortical Networks with Neuronal Avalanches. J. Neurosci. 32, 1061–1072 (2012).

45. McGinley, M. J., David, S. V. & McCormick, D. A. Cortical Membrane Potential Signature of Optimal States for Sensory Signal Detection. Neuron 87, 179–192 (2015).

46. Arieli, A., Sterkin, A. Grinvald, A. & Aertsen, A. Dynamics of ongoing activity: explanation of the large variability in evoked cortical responses. Science 273, 1868–1871 (1996).

47. Castro-Alamancos, M. A. Cortical Up and Activated States: Implications for Sensory Information Processing. Neurosci. 15, 625–634 (2009).

48. Azouz, R., & Gray, C. M. Cellular mechanisms contributing to response variability of cortical neurons in vivo. J. Neurosci. 19, 2209–2223 (1999).

49. Haider, B., Duque, A., Hasenstaub, A. R. Yu, Y. & McCormick, D. A. Enhancement of visual responsiveness by spontaneous local network activity in vivo. J. Neurophysiol. 97, 4186–202 (2007).

50. Gutnisky, D. A., Beaman, C. B. Lew, S. E. & Dragoi, V. Spontaneous Fluctuations in Visual Cortical Responses Influence Population Coding Accuracy. Cereb. Cortex bhv312 (2016). doi:10.1093/cercor/bhv312

51. Cowan, R. L. & Wilson, C. J. Spontaneous firing patterns and axonal projections of single corticostriatal neurons in the rat medial agranular cortex. J. Neurophysiol. 71, 17–32 (1994).

52. Paré, D., Shink, E. Gaudreau, H., Destexhe, A. & Lang, E. J. Impact of spontaneous synaptic activity on the resting properties of cat neocortical pyramidal neurons In vivo. J. Neurophysiol. 79, 1450–1460 (1998).

53. Markram, H., Wang, Y. & Tsodyks, M. Differential signaling via the same axon of neocortical pyramidal neurons. Proc Natl Acad Sci U S A 95, 5323–8. (1998).

54. Castro-Alamancos, M. A. & Connors, B. W. Distinct forms of short-term plasticity at excitatory synapses of hippocampus and neocortex. Proc. Natl. Acad. Sci. U. S. A. 94, 4161–6 (1997).

55. Castro-Alamancos, M. A. Cortical Up and Activated States: Implications for Sensory Information Processing. Neurosci. 15, 625–634 (2009).

56. Mancilla, J. G. & Ulinski, P. S. Role of GABA(A)-mediated inhibition in controlling the responses of regular spiking cells in turtle visual cortex. Vis. Neurosci. 18, 9–24 (2001).

57. Sachdev, R. N., Ebner, F. F. & Wilson, C. J. Effect of subthreshold up and down states on the whisker-evoked response in somatosensory cortex. J. Neurophysiol. 92, 3511–3521 (2004).

58. Civillico, E. F. & Contreras, D. Spatiotemporal properties of sensory responses in vivo are strongly dependent on network context. 6, 1–20 (2012).

59. Ollerenshaw, D. R. R., Zheng, H. J. V. J. V, Millard, D. C. C., Wang, Q. & Stanley, G. B. B. The adaptive trade-off between detection and discrimination in cortical representations and behavior. Neuron 81, 1152–1164 (2014).

60. Zheng, H. J. V., Wang, Q. & Stanley, G. B. Adaptive shaping of cortical response selectivity in the vibrissa pathway. J. Neurophysiol. 113, 3850–3865 (2015).

61. Shew, W. L. et al. Adaptation to sensory input tunes visual cortex to criticality. Nat. Phys. 11, 659–663 (2015).

62. Sekerli, M., Del Negro, C. A., Lee, R. H. & Butera, R. J. Estimating action potential thresholds from neuronal time-series: New metrics and evaluation of methodologies. IEEE Trans. Biomed. Eng. 51, 1665–1672 (2004).

